# The Multidimensional Battery of Prosody Perception (MBOPP)

**DOI:** 10.1101/555102

**Authors:** Kyle Jasmin, Frederic Dick, Adam Taylor Tierney

## Abstract

Prosody can be defined as the rhythm and intonation patterns spanning words, phrases and sentences. Accurate perception of prosody is an important component of many aspects of language processing, such as parsing grammatical structures, recognizing words, and determining where emphasis may be placed. Prosody perception is important for language acquisition and can be impaired in language-related developmental disorders. However, existing assessments of prosodic perception suffer from some shortcomings. These include being unsuitable for use with typically developing adults due to ceiling effects, or failing to allow the investigator to distinguish the unique contributions of individual acoustic features such as pitch and temporal cues. Here we present the Multi-Dimensional Battery of Prosody Perception (MBOPP), a novel tool for the assessment of prosody perception. It consists of two subtests – Linguistic Focus, which measures the ability to hear emphasis or sentential stress, and Phrase Boundaries, which measures the ability to hear where in a compound sentence one phrase ends, and another begins. Perception of individual acoustic dimensions (Pitch and Time) can be examined separately, and test difficulty can be precisely calibrated by the the experimenter because stimuli were created using a continuous voice morph space. We present validation analyses from a sample of 57 individuals and discuss how the battery might be deployed to examine perception of prosody in various populations.

## 1. Introduction

### 1.1 Multiple Dimensions for Prosody

One of the main tasks in speech perception is thought to be categorizing rapidly-evolving speech sounds into linguistically informative phonemes or syllables. However, speech contains acoustic patterns on slower time scales as well. These suprasegmental or *prosodic* patterns convey crucial disambiguating lexical, syntactic, and emotional cues that help the listener capture the intended message of the talker. In English, prosodic features can be conveyed by many acoustic dimensions, including changes in pitch, amplitude, and the duration of elements. For example, prosodic *focus*, which helps listeners direct attention to particularly important words or phrases in a sentence, is typically cued by an increase in the amplitude and duration of the emphasized elements, along with exaggerated pitch excursion (Breen, Kaswer, Van Dyke, Krivokapic, & Landi, 2010; see Figure 1a-b for an example). Listeners can use focus to determine the portion of the sentence to which they should be directing their attention. Similarly, *lexical stress* is cued by a combination of increased amplitude, pitch changes, and increased syllable duration (Chrabaszcz, Winn, Lin, & Idsardi,, 2014; Mattys, 2000). Listeners can use stress to help distinguish between different words (i.e. “PREsent” versus “preSENT”) and to detect word boundaries (Nakatani & Schaffer, 1977). Finally, *phrase boundaries* tend to coincide with a change in pitch and lengthening of the syllable just prior to the boundary (Choi, Haswegawa, & Cole, 2005; Cumming, 2010; de Pijper & Sanderman, 1994; Streeter, 1978).

**Figure 1:**
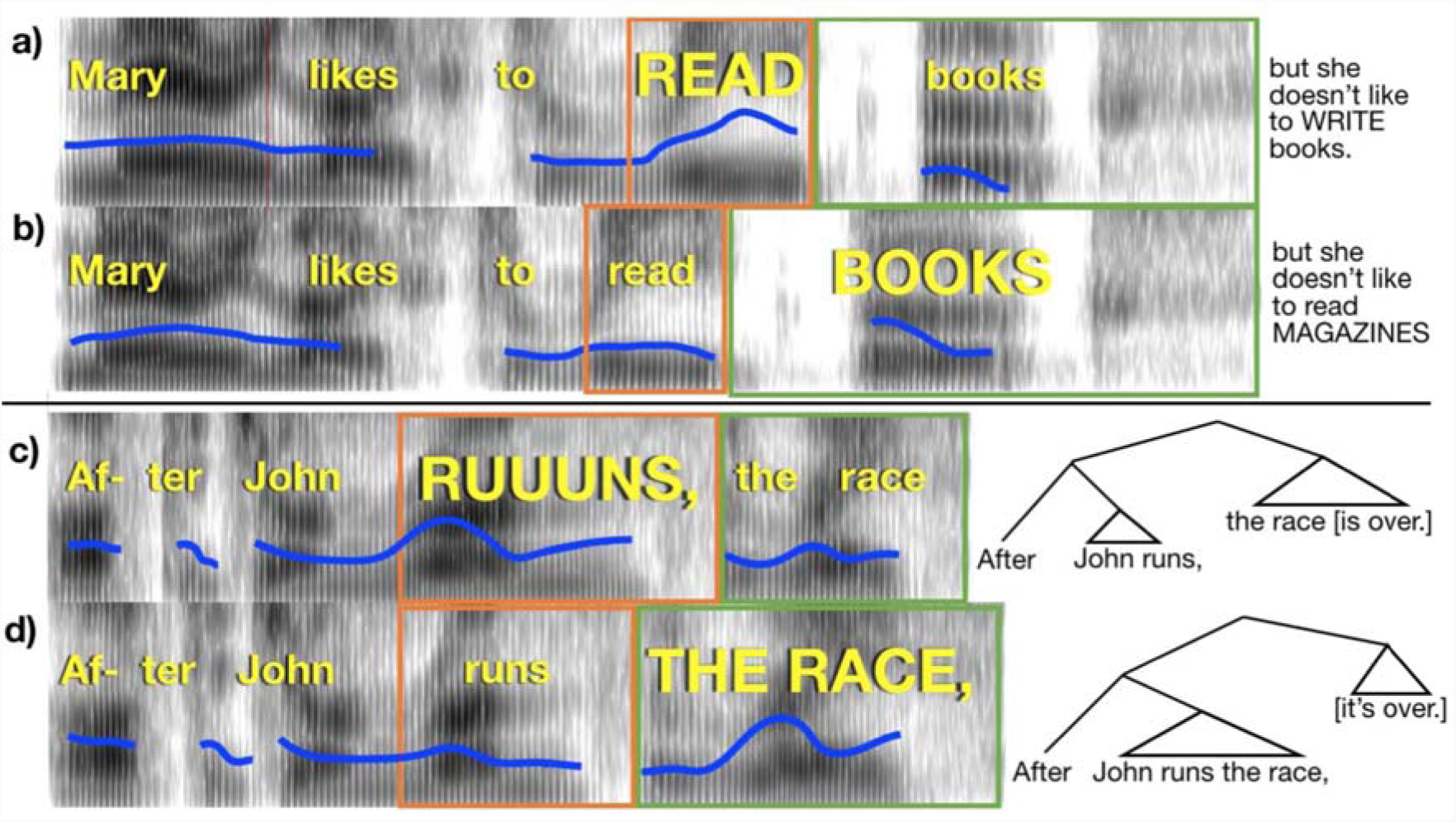
Pitch and duration (time) correlates of emphatic accents and phrase boundaries. Example spectrograms of stimuli used in the experiment (time on horizontal axis, frequency on vertical axis, and amplitude in grayscale), with linguistic features cued simultaneously by pitch and duration (the “Combined” condition). Blue line indicates the fundamental frequency of the voice. Width of orange and green boxes indicate duration of the words within the box. A) Emphatic accent places focus on “read”. Completion of the sentence appears to the right. B) Emphatic accent places focus on “books”; sentence completion is at right. C) A phrase boundary occurs after “runs”. D) A phrase boundary occurs after “race”. Syntactic trees are indicated at right to illustrate the structure conveyed by the acoustics of the stimuli.

Listeners can make use of such prosodic cues to clarify potentially ambiguous syntactic structures in a sentence (Beach, 1991; Frazier, Carlson, & Clifton Jr, 2006; Lehiste, Olive, & Streeter, 1976; Marslen-Wilson, Tyler, Warren, Grenier, & Lee,, 1992; Jasmin, Dick, Holt, & Tierney, 2018). In fact, prosodic patterns may be a more powerful cue to phrase structure than statistical patterns, as artificial grammar learning experiments have shown that when prosodic cues and transitional probabilities are pitted against one another, listeners will learn hierarchical structure which reflects prosodic information (Langus, Marchetto, Bion, & Nespor, 2012).

### 1.2 Prosody and Language Acquisition

Given the useful information prosodic cues provide about the structure of language, accurate prosody perception may be a crucial foundational skill for successful acquisition of language. Indeed, phonemic and prosodic awareness are independent predictors of word reading (Clin, Wade-Woolley, & Heggie, 2009; Holliman, Wood, & Sheehy, 2010a; Defior, Gutierrez-Palma, & Cano-Marin, 2012; Goswami et al., 2013; Jimenez-Fernandez, Gutierrez-Palma, & Defior, 2015; Wade-Woolley, 2016; for a review see Wade-Woolley & Heggie, 2015), suggesting that prosody perception forms a separate dimension of linguistic skill relevant to reading acquisition. Not only has dyslexia has been linked to impaired prosody perception (Goswami, Gerson, & Astruc, 2010; Holliman et al., 2010a; Mundy & Carroll, 2012; Wade-Woolley, 2016; Wood & Terrell, 1998), but in adolescents with dyslexia, difficulties with the perception of lexical stress have been shown to be more prominent than problems with segmental phonology (Anastasiou & Protopapas, 2014). Finally, prosodic sensitivity also predicts word reading one year later (Holliman, Wood, & Sheehy, 2010b; Calet, Gutierrez-Palma, Simipson, Gonzalez-Trujillo, & Defior, 2015), suggesting that prosody perception is a foundational skill upon which children draw when learning to read.

Such links between prosodic awareness and language acquisition suggest that the difficulties with prosody perception that accompany certain clinical diagnoses may have consequences for language acquisition. For example, some individuals with autism spectrum disorders (ASD) produce speech which lacks the usual acoustic characteristics which mark particular prosodic features; for example, the difference in duration between stressed and unstressed syllables tends to be smaller in the speech of children with ASD (Paul, Bianchi, Augustyn, Klin, & Volkmar, 2008). These prosodic production deficits extend to perception as well: individuals with ASD tend to have difficulty with the perception of prosodic cues to emotion (Globerson, Amir, Kishon-Rabin, & Golan, 2015; Golan, Baron-Cohen, Hill, & Rutherford, 2007; Kleinman, Marciano, & Ault, 2001; Phillip et al., 2010; Rutherford, Baron-Cohen, & Wheelwright, 2002), lexical stress (Kargas, López, Morris, & Reddy, 2016), phrase boundaries (Diehl, Bennetto, Watson, Gunlogson, & McDonough, 2008), and linguistic focus (Peppé, Cleland, Gibbon, O’Hare, & Castilla, 2011) in speech (but see Diehl, Friedberg, Paul, & Snedeker, 2015). These prosody perception difficulties can interfere not only with communication skill and sociability (Paul, Augustyn, Klin, & Volkmar, 2005), but may also increase the risk of delayed language acquisition given the importance of prosody for disambiguating language meaning (Lyons, Simmons, & Streeter, 2014).

### 1.3 Prosody and Language Disorders

Prosody perception is, therefore, a vital skill supporting language development, and is impaired in several clinical populations in which there is intense interest. As mentioned above, prosodic features tend to be conveyed by a mixture of multiple different cues, including changes in the pitch and duration of syllables and words. As a result, one source of difficulties with prosody perception may be impairments in auditory processing, a possibility supported by findings that prosody perception in children correlates with psychophysical thresholds for pitch, duration, and amplitude rise time (Goswami et al., 2013; Haake et al., 2013; Richards & Goswami, 2015). However, impairments in auditory processing can be present for one dimension in the presence of preserved processing in other dimensions. In particular, impaired pitch perception can co-occur with preserved duration perception (and vice versa – Kidd, Watson, & Gygi, 2007). Similarly, research on amusia has shown that highly impaired memory for pitch sequences can co-occur with preserved memory for durational sequences (Hyde & Peretz, 2004). A prosody perception deficit in a given individual, therefore, could reflect impaired pitch perception or duration perception or both. Existing methodologies for assessing prosody perception, however, cannot control the acoustic cues to different prosodic features, and therefore cannot diagnose the source of an individual’s prosodic impairment.

### 1.4 Existing Prosody Tests

Although there exist many widely available standardized tests of segmental speech perception usable by individuals of all ages (Killion, Niquette, Gudmundsen, Revit, & Banerjee, 2004; Nilsson, Soli, & Sullivan, 1994; Wilson, 2003), there are comparatively few instruments publicly available for researchers and clinicians interested in testing suprasegmental speech perception. As a consequence, prosody perception research has been carried out using a wide variety of in-house methods developed within single laboratories, making comparison across studies difficult. These include perceptual matching tasks such as matching low-pass filtered sentences or indicating whether the prosodic structure of low-pass filtered sentences match unfiltered target sentences (Cumming, Wilson, Leong, Colling, & Goswami, 2015; Fisher, Plante, Vance, Gerken, & Glattke, 2007; Wood & Terrell, 1998). Participants have also been asked to match the stress pattern of a nonsense phrase like “DEEdee DEEdee” with a spoken target phrase like “Harry Potter” (Goswami et al., 2010; Holliman, Wood, & Sheehy, 2012; Mundy & Carroll, 2012; Whalley & Hansen, 2006). These tests have the advantage of isolating the suprasegmental elements of speech. However, these tests do not require prosodic categorization, and therefore arguably measure auditory discrimination rather than prosody perception per se. Moreover, these tests are not publicly available.

The most widely used battery of prosody perception available for purchase by the public is the Profiling Elements of Prosodic Systems—Children test, or PEPS-C (Peppé & McCann, 2003). This test assesses the perception and production of four different aspects of prosody: affect, phrase structure, focus, and interaction. Each subtest features two different sets of trials. In “form” trials, the listener is asked to make same/different judgments on utterances which either do or do not differ based on a prosodic feature. In “function” trials, the listener is asked to infer the speaker’s intent by detecting a prosodic feature. For example, one item from the phrase structure subtest asks listeners to point to the picture that best fits the utterance “fish, fingers, and fruit” (as opposed to “fish fingers and fruit”; NB:British English “fish fingers” are called “fish sticks” in American English). This test has been successfully used to study a variety of topics related to prosody perception in children, including the relationship between prosody perception and reading ability in typically developing children (Lochrin, Arciuli, & Sharma, 2015), and impairments in prosody perception in children with specific language impairment, dyslexia, and ASD (Wells & Pepé, 2003; Jarvinen-Pasley, Peppé, King-Smith, & Heaton, 2008; Marshall, Harcourt-Brown, Ramus, & van der Lely, 2009).

The main limitation of the PEPS-C is that it was designed to be administered to children, and therefore many adults would perform at ceiling. The PEPS-C was adapted from an earlier battery designed to be used with adults (the PEPS), but it is not available for use by the public, and there is also evidence for the existence of ceiling effects in adult PEPS data (Peppé, Maxim, & Wells, 2000). Moreover, there are a number of examples of ceiling effects in the literature on prosody perception in adolescents and adults in research using other prosody perception tests (Chevallier, Noveck, Happé, & Wilson, 2008; Lyons et al., 2014; Paul et al., 2005), suggesting that existing methodologies for testing prosody perception are insufficiently challenging for adult participants. Research on prosody would be facilitated by a publicly available test with adaptive difficulty suitable for a range of ages and backgrounds.

### 1.5 The Current Study

Here we report and make publicly available the Multidimensional Battery of Prosody Perception (MBOPP), a battery of prosody perception with adaptive difficulty which is therefore suitable for participants of all ages, backgrounds, and ability levels. This battery consists of two tests, one assessing the perception of linguistic focus and another assessing the perception of phrase boundaries. For both tests, stimuli were constructed by asking an actor to read aloud sequences of words which were identical lexically but differed on the presence of a prosodic feature. Thus, each sentence in the focus test has an “early focus” and “late focus” version, referring to the relative position of emphasized elements. Similarly, the sentences in the phrase test have an “early closure” and “late closure” version, referring to the placement of the phrase boundary (indicated typographically with a comma). Speech morphing software (STRAIGHT, Kawahara & Irino, 2005) was then used to morph these two recordings onto one another, such that the extent to which pitch and durational patterns cued the existence of one versus the other prosodic interpretation could be varied independently. This method allows the researcher to tune the difficulty of the test to any population (by choosing which subset of stimuli to use), and also enables investigation of cue-specific prosody perception. This test was presented to 57 typically developed adult participants to examine the relative usefulness of pitch versus durational cues for focus and phrase boundary perception, and to measure the reliability of each subtest.

## 2. Methods

### 2.1 Participants

Participants (N=57, 29F, 28M, aged 34.4±12.8) were recruited using Birkbeck’s SONA system in exchange for payment. The same participants completed both the focus perception and phrase perception tasks.

### 2.2 Materials – Focus Perception

The Focus Perception test consists of 47 compound sentences (two independent clauses separated by a conjunction; Table 1). We recorded spoken versions of these sentences in a quiet room using a Rode NT1-A condenser microphone (44.1 kHz, 32-bit) as they were spoken by a former professional actor, now a speech researcher. The actor placed contrastive accents to emphasize the capitalized words in the sentences. Each of the sentences was read with emphasis on two different word pairs, thus creating two versions: an “early focus” version (e.g., “<u>Mary likes to READ books</u>, but she doesn’t like to WRITE them,” focus indicated by upper-case letters), and “late focus”, where the focus elements occurred in later positions in the sentence (e.g., “<u>Mary likes to read BOOKS</u>, but she doesn’t like to read MAGAZINES,” focus indicated by upper-case letters; Figure 1a-b). Thus the emphasis placed on the words in capitalized letters served to indicate *contrastive focus,* meant to indicate which linguistic elements (words, in this case) should receive greater attention in order to clarify the speaker’s intentions. For example, suppose the conversation began as follows:

**Table 1.**
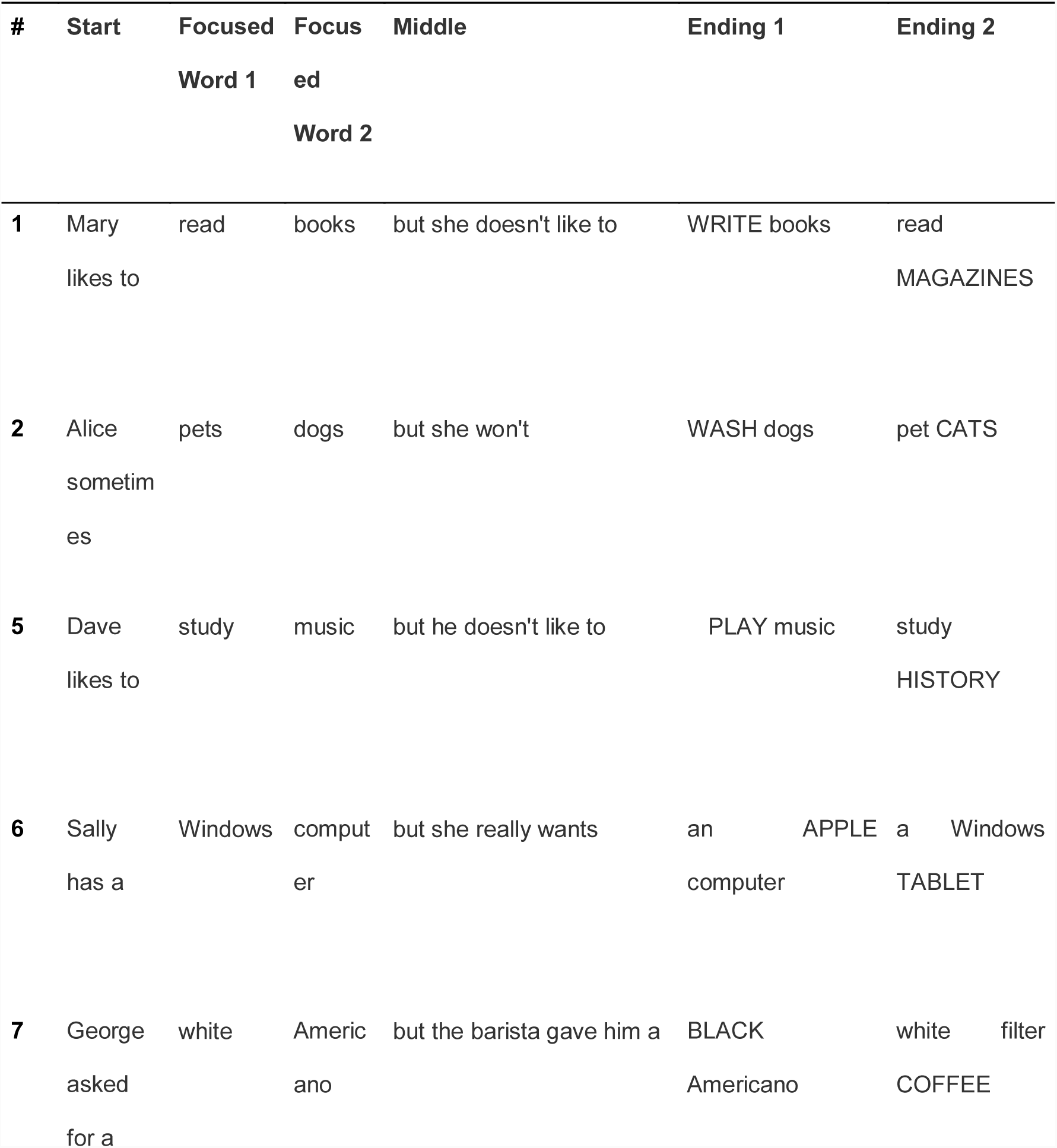
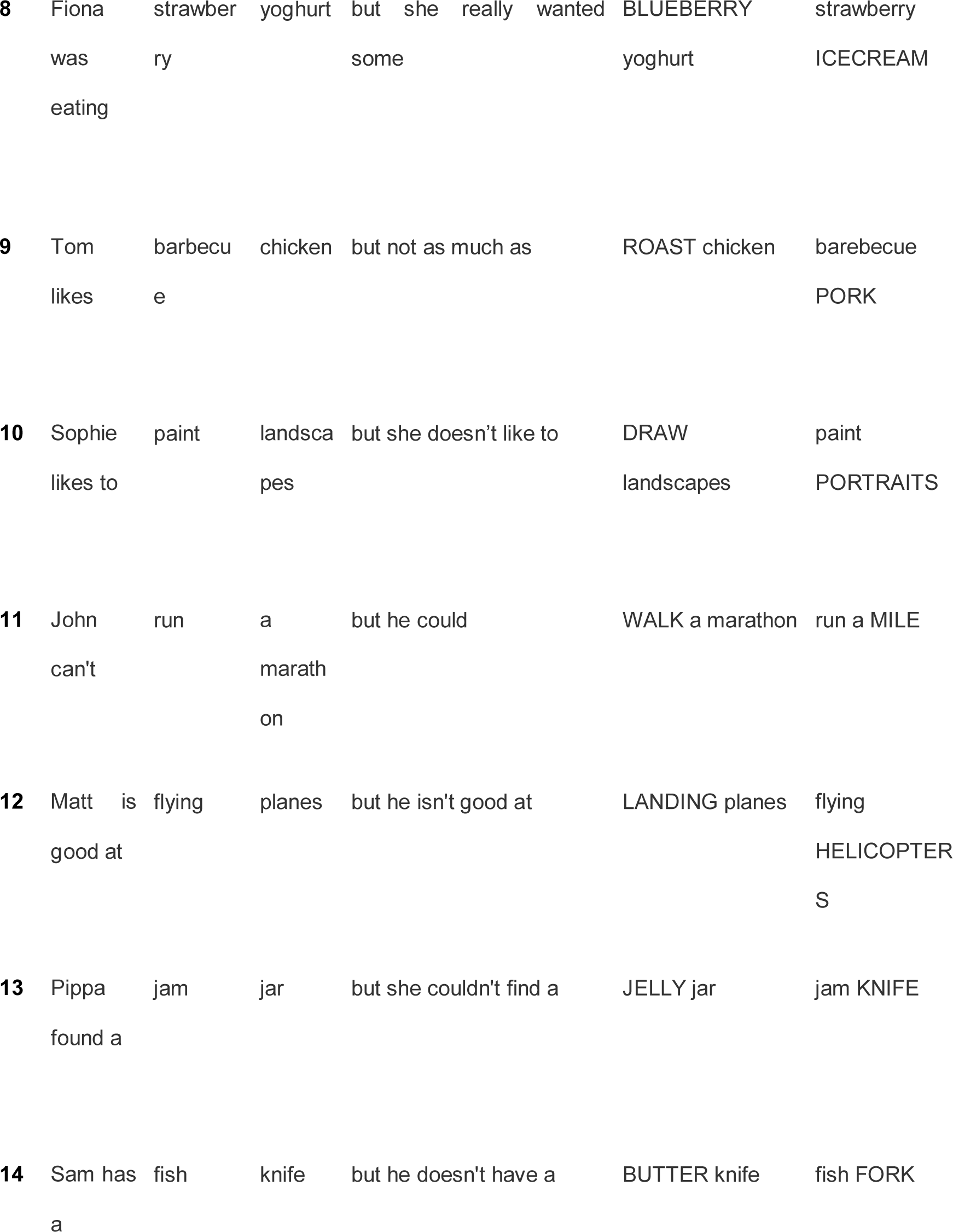
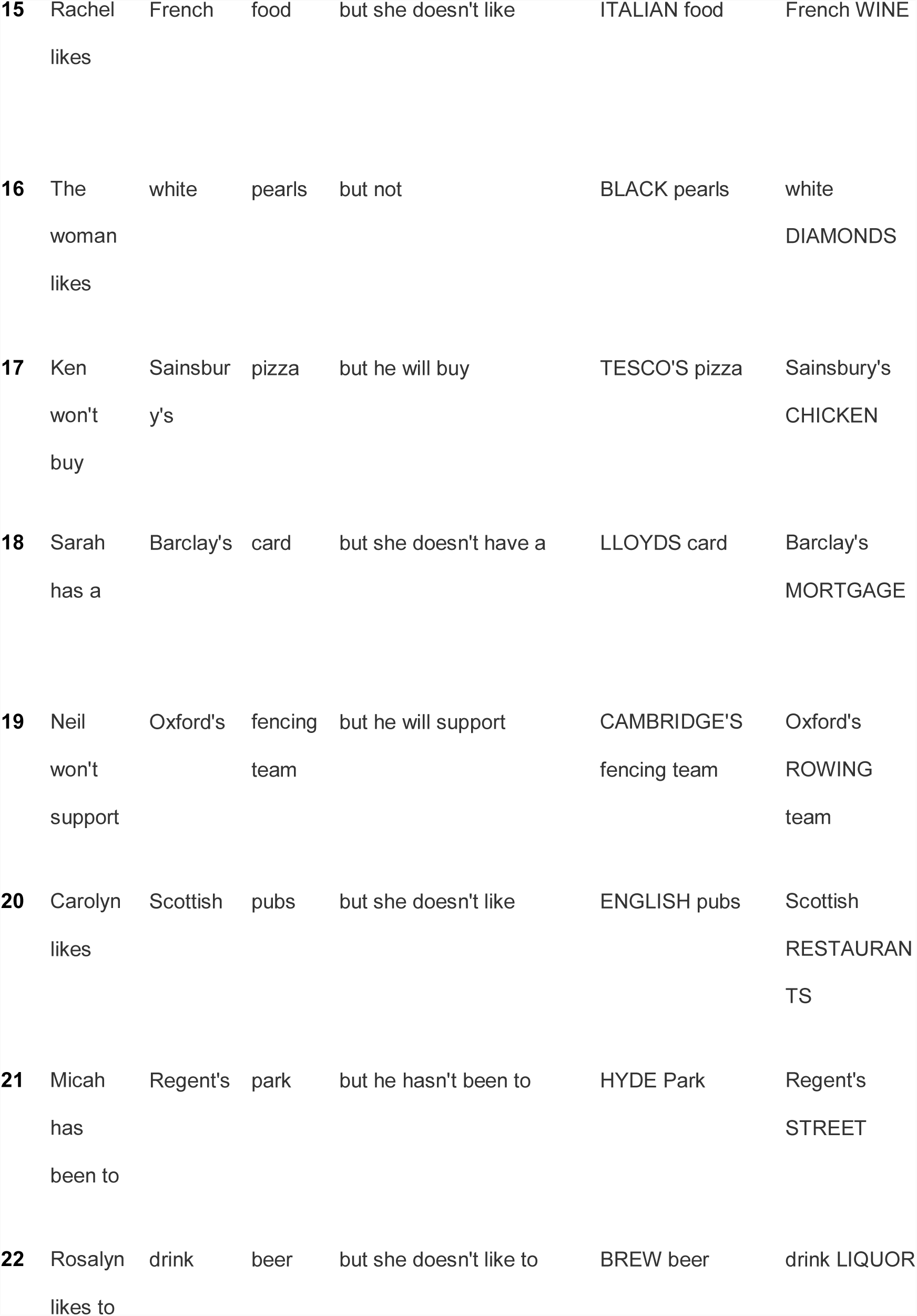
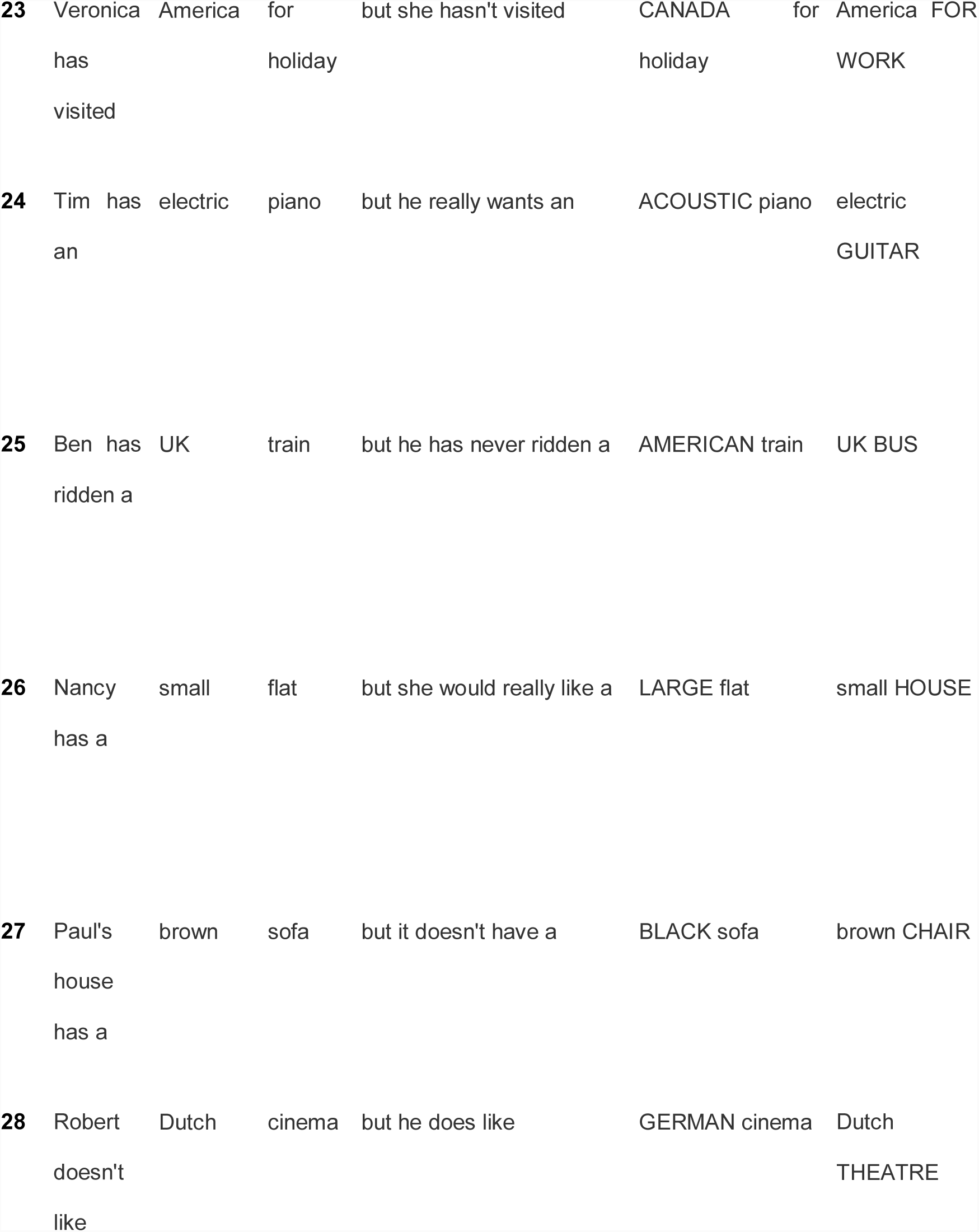
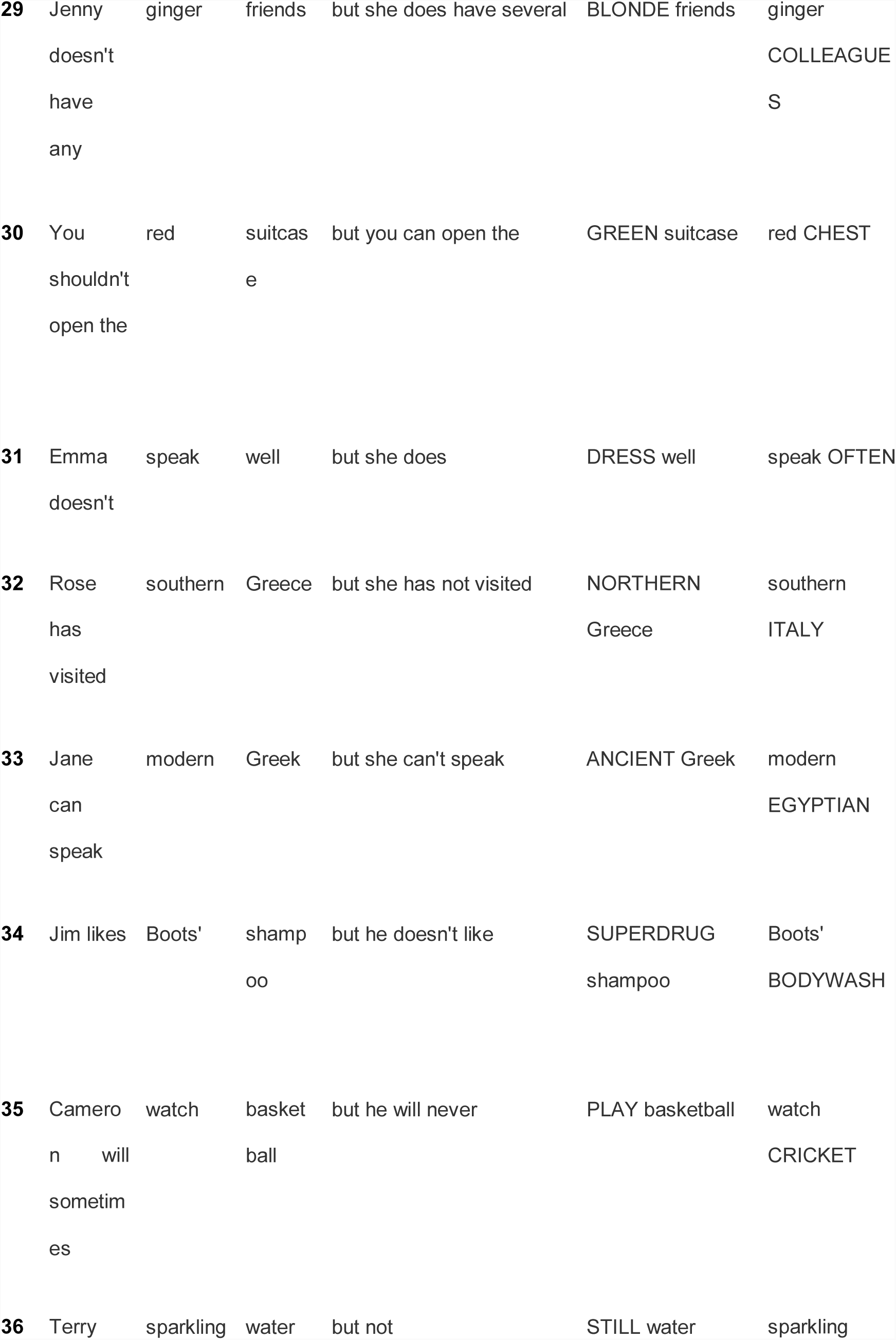
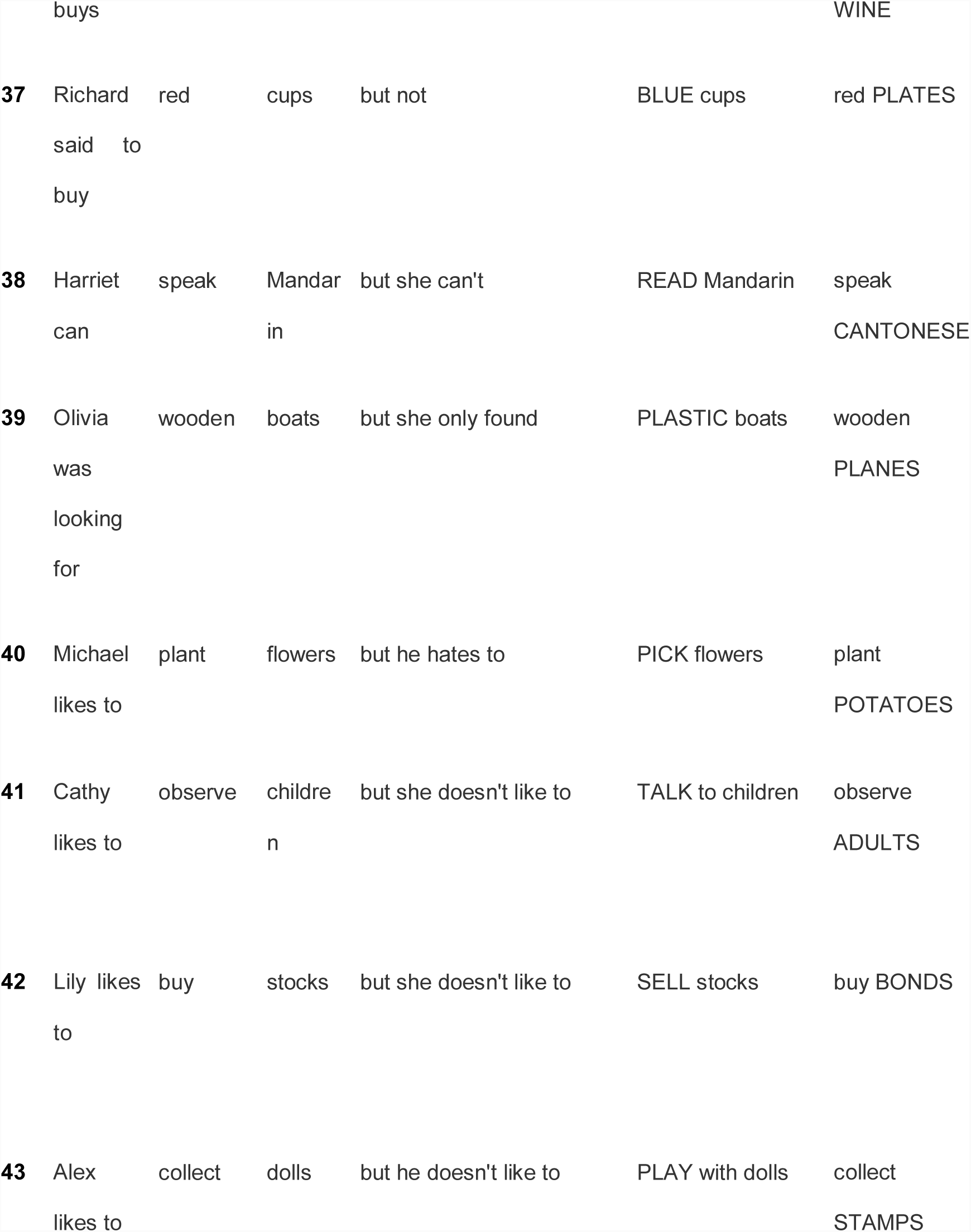
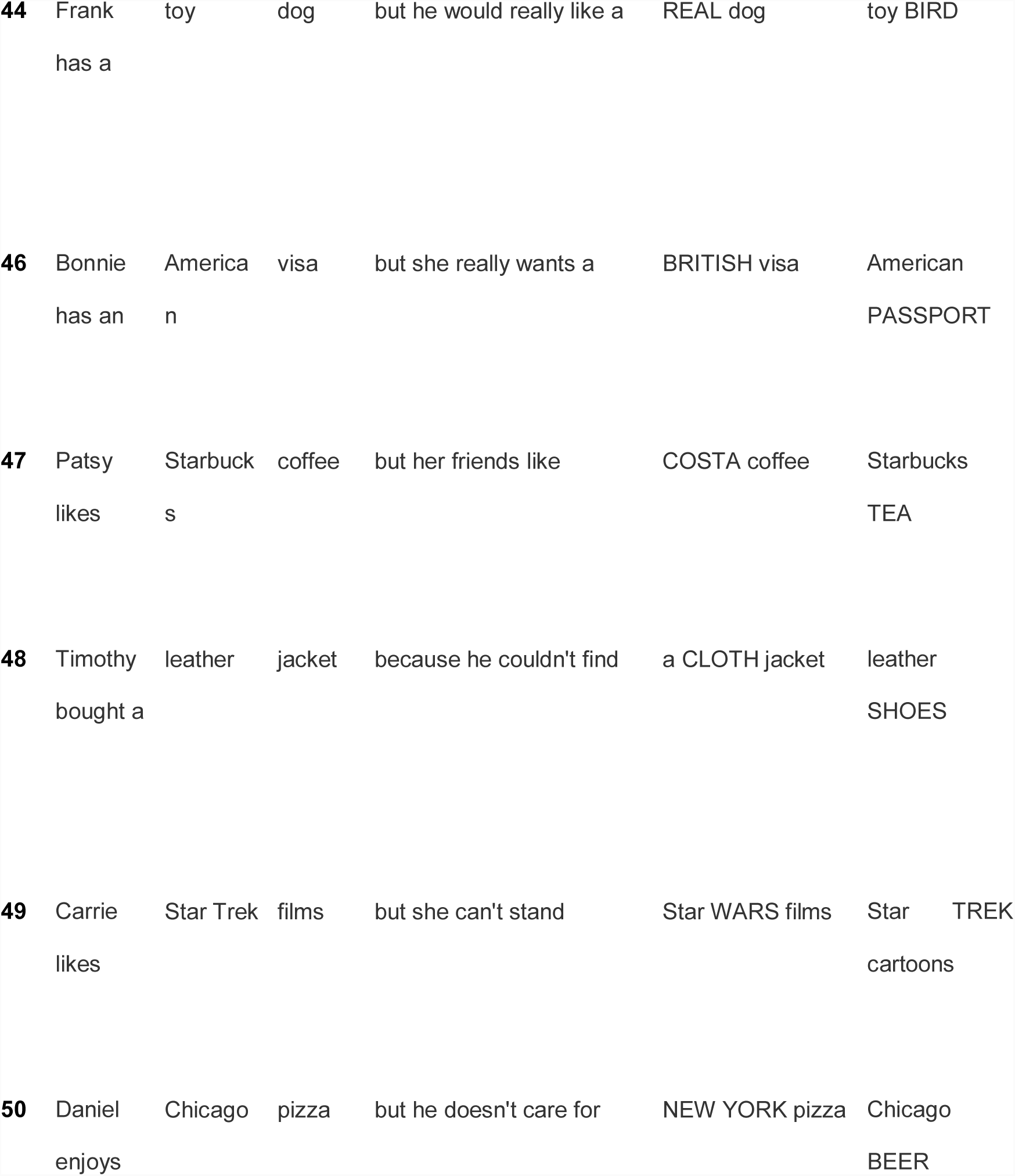
Text of Focus Stimuli Sentences

A. Why doesn’t Mary like books?
B. She likes to READ books, but not WRITE them.

The focused elements spoken by B serve to contrast with the presupposition by speaker A. The terms “early focus” and “late focus” used in this article refer simply to which pair of words is emphasized (e.g. READ and WRITE occur earlier than BOOKS and MAGAZINES, respectively.)

The audio recordings of these sentences were trimmed such that they included only the first clause, which consisted of identical words in each version (this clause is indicated in the examples above via underlining). The raw recordings of “early” and “late” focus sentences were then morphed together to create intermediate versions. Morphing was performed with STRAIGHT software (Kawahara & Irino, 2005). The two recordings of each sentence (differing only in the placement of the emphasized word) were manually time-aligned by marking the onsets and offsets of the corresponding phonemes in each recording. After establishing these corresponding ‘anchor points’, morphed intermediate versions of the sentences were synthesized. An experimenter listened to the result of the morphing in order to check the quality of the output. If quality was low, anchor points were added or adjusted and the procedure was repeated, until the resulting morph sounded natural. STRAIGHT allows morphs along several dimensions: Aperiodicity, Spectrum, Frequency, Time (duration), and F0 (pitch). For the morphs created for this prosody battery, only Duration and Pitch were manipulated.

We are distributing this stimulus set with morphs in three conditions: Pitch-Only, Time-Only, and Combined. The Combined condition consists of stimuli in which duration and pitch information cue emphasis on the same word – either early focus or late focus (e.g. Mary likes to READ books vs Mary likes to read BOOKS). Morphing rates are expressed in terms of percent, such that lower values indicate more information from the early focus recording, and higher values indicate more information from the late focus recording, while 50% indicates an equal amount of a given dimension from each recording.

For stimuli in the Pitch-Only condition, the emphasized word in the sentence is conveyed by pitch cues alone which vary from 0% (pitch information coming entirely from the early focus recording) to 100% (pitch information coming from the late focus recording), while duration cues are ambiguous with the Time parameter always set at 50%. In the Time-Only condition, emphasis is conveyed only by durational cues, which similarly vary from 0% to 100%, while pitch cues are ambiguous, always set at 50%. The other morphing dimensions available in STRAIGHT (Aperiodicity, Spectrum, and Frequency) were held at 50% such that morphs contained equal amounts of information from the two recordings.

Table 2 displays the morphings rates included in the stimuli published with this article. The filenames format for the stimuli is as follows.

**Table 2.** Morphing rates for Phrase and Focus test stimuli.

| Condition | Pitch Morphing Rate | Duration Morphing Rate |
| --- | --- | --- |
| Pitch-Only | 0% to 40%, 60 to 100%, in 5% increments | Always 50% |
| TimeOnly | Always 50% | 0% to 40%, 60 to 100%, in 5% increments |
| Combined | 0% to 40%, 60 to 100%, in 5% increments | 0% to 40%, 60 to 100%, in 5% increments |

[Stimulus number] _ [pitch morphing rate] _ [time morphing rate] .wav

Examples:

Focus1_pitch0_time0.wav – pitch and duration both cue EARLY focus (Combined)

Focus1_Pitch100_time100.wav – pitch and duration both cue LATE focus (Combined)

Focus1_pitch50_time0.wav – pitch is ambiguous, only duration cues EARLY focus (Time-Only)

Focus1_pitch50_time100.wav – pitch is ambiguous, only duration cues LATE focus (Time-Only)

Focus1_pitch0_time50.wav – duration is ambiguous, only pitch cues EARLY focus (Pitch-Only)

Focus1_pitch100_time50.wav – duration is ambiguous, only pitch cues LATE focus (Pitch-Only)

For the experiments included in this report, these 6 different kinds of morphs were created by varying the amount of pitch-related and time information either independently or simultaneously. For the Pitch-Only condition, duration morphing rates were held at 50%, while two contrasting pitch versions were created at 25% (towards early focus) and 75% (towards late focus). For the Time-Only condition, pitch was held at 50% while duration was manipulated to be 25% (early focus) or 75% (late focus). For the Combined condition, both the pitch and the Duration dimensions were manipulated simultaneously to be 25% or 75%. Morphing rates of 25% (instead of 0%) and 75% (instead of 100%) were used to make the task more difficult. The task could be made more difficult by moving these values even closer to 50% (e.g. 40% for early focus and 60% for late focus). All files were saved and subsequently presented at a sampling rate 44.1 kHz with 16-bit quantization.

The text of the stimuli are given in Table 1. The auditory recordings consist of the following portions of the text: Start, Focused Word 1, Focused Word 2.

### 2.3 Procedure – Focus perception

Performance and reliability data reported here were collected with Psychtoolbox in Matlab. We tested participants’ ability to detect prosodic differences by asking them to match auditory versions of sentences with text ones. Participants read sentences presented visually on the screen one at a time, which were either early or late focus. For example, one visually presented sentence was “Mary likes to READ books, but she doesn’t like to WRITE books.”

The emphasized words appeared in all upper-case letters, as in the example above. Subjects were then given 4 seconds to read the sentence to themselves silently and imagine how it should sound if someone spoke it aloud. Following this, subjects heard the early focus and late focus versions of the first independent clause of the stimulus sentence (up to but not including the conjunction). The order of the presentation was randomized. Participants decided which of the two readings contained emphasis placed on the same word as in the text sentence and responded by pressing “1” or “2” on the keyboard to indicate if they thought the first version or second version was spoken in a way that better matched the on-screen version of the sentence. The stimuli were divided into 3 lists (47 trials each) and counterbalanced such that participants heard an equal number of Pitch-Only, Time-Only and Combined stimulus examples. For half (23) of the stimuli, two of the presentations were early focus, and one was late focus; for the remaining stimuli, two presentations were late focus and one was early. The entire task lasted approximately 30 minutes.

### 2.4 Materials – Phrase Perception

The Phrase Perception test stimuli consisted of 42 pairs of short sentences with a subordinate clause appearing before a main clause (see Figure 1c-d). About half of these came from a published study (Kjelgaard & Speer, 1999) and the rest were created for this test (see Table 3). The sentence pairs consisted of two similar sentences, the first several words of which were identical. In the first type of sentence, “early closure”, the subordinate clause’s verb was used intransitively, and the following noun was the subject of a new clause (“After John runs, the race is over”). In the second type of sentence, “late closure”, the verb was used transitively and took the immediately following noun as its object, which caused a phrase boundary to occur slightly later in the sentence than in the early close version (“After John runs the race, it’s over”). Both versions of the sentence were lexically identical from the start of the sentence until the end of the second noun. The same actor recorded early and late closure versions of the sentences in his own standard Southern English dialect. The recordings were cropped such that only the lexically identical portions of the two versions remained, and silent pauses after phrase breaks were removed.

**Table 3.** Text of the Phrase Test sentences, each of which has two versions, where a phrase boundary occurs either earlier or later in the sentence.

| # | Closure | Start | Finish |
| --- | --- | --- | --- |
| 1 | Early | After Jane dusts, the dining table | is clean |
| 1 | Late | After Jane dusts the dining table, | it's clean |
| 2 | Early | After John runs, the race | is over |
| 2 | Late | After John runs the race, | it's over |
| **5** | Early | Because Mike phoned, his mother | was relieved |
| **5** | Late | Because Mike phoned his mother, | she was relieved |
| **7** | Early | Because Sarah answered, the teacher | was proud |
| **7** | Late | Because Sarah answered the teacher, | she was proud |
| **8** | Early | Because Tara cleaned, the house | was spotless |
| **8** | Late | Because Tara cleaned the house, | it was spotless |
| **9** | Early | Because George forgot, the party | had started |
| **9** | Late | Because George forgot the party, | he was sad |
| **10** | Early | Because Mike paid, the bill | was smaller |
| **10** | Late | Because Mike paid the bill, | it was smaller |
| **13** | Early | If Charles is baby-sitting, the children | are happy |
| **13** | Late | If Charles is baby-sitting the children, | they're happy |
| **14** | Early | If George is programming, the computer | is busy |
| **14** | Late | If George is programming the computer, | it's busy |
| **15** | Early | If Ian doesn't notice, Beth | is fine |
| **15** | Late | If Ian doesn't notice Beth, | it's fine |
| **16** | Early | If Joe starts, the meeting | will be long |
| **16** | Late | If Joe starts the meeting, | it'll be long |
| **18** | Early | If Laura is folding, the towels | will be neat |
| **18** | Late | If Laura is folding the towels, | they'll be neat |
| **19** | Early | When the baby finishes, the bottle | will be empty |
| **19** | Late | When the baby finishes the bottle, | it'll be empty |
| **20** | Early | If Barbara gives up, the ship | will be plundered |
| **20** | Late | If Barbara gives up the ship, | it'll be plundered |
| **21** | Early | If the Scissor Sisters open, the show | will be great |
| **21** | Late | If the Scissor Sisters open the show, | it'll be great |
| **22** | Early | If the maid packs, the suitcase | will be tidy |
| **22** | Late | If the maid packs the suitcase, | it'll be tidy |
| **23** | Early | If Tom wins, the contest | is over |
| **23** | Late | If Tom wins the contest, | it's over |
| **24** | Early | If the doctor calls, your sister | will answer |
| **24** | Late | If the doctor calls your sister, | she'll answer |
| **25** | Early | If Jack cleans, the kitchen | will be filthy |
| **25** | Late | If Jack cleans the kitchen, | it'll be filthy |
| **26** | Early | If dad digs, the hole | will be deep |
| **26** | Late | If dad digs the hole, | it'll be deep |
| **27** | Early | When a man cheats, his friends | get angry |
| **27** | Late | When a man cheats his friends, | they're angry |
| **29** | Early | When Gaga sings, the song | is a hit |
| **29** | Late | When Gaga sings the song, | it's a hit |
| **30** | Early | When Roger leaves, the house | is dark |
| **30** | Late | When Roger leaves the house, | it's dark |
| **31** | Early | When Suzie visits, her grandpa | is happy |
| **31** | Late | When Suzie visits her grandpa, | he's happy |
| **32** | Early | When the clock strikes, the hour | has started |
| **32** | Late | When the clock strikes the hour, | it's started |
| **33** | Early | When the guerrillas fight, the battle | has begun |
| **33** | Late | When the guerrillas fight the battle, | it's begun |
| **34** | Early | When the maid cleans, the rooms | are organized |
| **34** | Late | When the maid cleans the rooms, | they're organized |
| **35** | Early | When the original cast performs, the play | is fantastic |
| **35** | Late | When the original cast performs the play, | it's fantastic |
| **36** | Early | When Tim is presenting, the lectures | are interesting |
| **36** | Late | When Tim is presenting the lectures, | they're interesting |
| **37** | Early | When The Beatles play, the music | is noisy |
| **37** | Late | When The Beatles play the music, | it's noisy |
| **38** | Early | When Paul drinks, the rum | disappears |
| **38** | Late | When Paul drinks the rum, | it disappears |
| **39** | Early | When Mary helps, the homeless | are grateful |
| **39** | Late | When Mary helps the homeless, | they're grateful |
| **40** | Early | When the phone loads, the app | crashes |
| **40** | Late | When the phone loads the app, | it crashes |
| **41** | Early | When the shop closes, its doors | are locked |
| **41** | Late | When the shop closes its doors, | they're locked |
| **42** | Early | When a train passes, the station | shakes |
| **42** | Late | When a train passes the station, | it shakes |
| **43** | Early | When the actor practices, the monologue | is excellent |
| **43** | Late | When the actor practices the monologue, | it's excellent |
| **44** | Early | When the cowboy rides, the horse | is tired |
| **44** | Late | When the cowboy rides the horse, | it's tired |
| **46** | Early | Whenever the guard checks, the door | is locked |
| **46** | Late | Whenever the guard checks the door, | it's locked |
| **47** | Early | Whenever Bill teaches, the course | is boring |
| **47** | Late | Whenever Bill teaches the course, | it's boring |
| **48** | Early | Whenever a customer tips, the waiter | is pleased |
| **48** | Late | Whenever a customer tips the waiter, | he's pleased |
| **49** | Early | Whenever Rachel leads, the discussion | is exciting |
| **49** | Late | Whenever Rachel leads the discussion, | it's exciting |
| **50** | Early | Whenever Mary writes, the paper | is excellent |
| **50** | Late | Whenever Mary writes the paper, | it's excellent |

Auditory stimuli for the phrase test were created in the same way as in the focus test, by asking an actor to read aloud the two versions of each sentence (the early and late closure). Then the recordings were cropped to the lexically identical portions, corresponding anchor points were defined, and morphs were created in STRAIGHT. The morphs we publish here were created with the same proportions as in the focus test (Table 2).

#### Phrase Perception Test Procedure

For the validation experiments reported here, we used stimuli with early or late closure cued by 75% and 25% morphing rates. The procedure for the Linguistic Phrase test was similar to that of the Linguistic Focus Test. On each trial, participants read a text version of each sentence online, which was either early or late closure, as indicated by the grammar of the sentence and a comma placed after the first clause (Figure 1c-d). Participants read the sentence to themselves silently and imagined how it should sound if someone spoke it aloud. Following this, subjects heard the first part of the sentence (which was identical in the early and late closure versions) spoken aloud, in two different ways, one that cued an early closure reading and another that cued a late closure reading. Participants decided which of the two readings best reflected the text sentence (and the location of its phrase boundary, indicated grammatically and orthographically with a comma) and responded by pressing “1” or “2” on the keyboard to indicate if they thought the first version or second version was spoken in a way that better matched the on-screen version of the sentence. The grammatical difference between the two spoken utterances on each trial was cued by pitch differences (Pitch-Only), duration differences (Time-Only), or both pitch and duration differences (Combined). Subjects completed three blocks of 42 trials. Stimuli were counterbalanced and half of the presentations were early close and half were late close. The task was performed in a lab at Birkbeck and lasted approximately 25 minutes.

## 3. Results

### 3.1 Overall performance

Figures 2 and 3 display all participants’ performance in the phrase perception and focus perception tests, respectively. Performance across participants is summarized in Table 4, which describes performance across deciles for each test. Although there was overall a very wide range of performance, the extent to which ceiling effects were present varied across the subtests. For the phrase perception subtests, ceiling performance was not evident: greater than 95% performance was achieved by less than 10% of participants for the Pitch-Only condition and Time-Only conditions and by only the top 20% for the both condition. Nevertheless, floor effects were also not evident, with less than 55% performance achieved by only the bottom 20% for the Pitch-Only condition and the bottom 10% for the Time-Only and both conditions. The focus perception subtests, on the other hand, showed more evidence of ceiling effects.. In the Pitch-Only condition, 40% of participants achieved greater than 95% performance. In the Combined condition, this rose to more than 50% of participants. Less than 10% of participants, however, achieved this score in the Time-Only condition. These results suggest that to avoid ceiling effects in typically developing adults, cue magnitude for the focus test should be decreased slightly. Given these ceiling effects, rau transforms were applied to all data prior to further analysis. There was no indication of floor effects – near-chance scores (less than 55%) were achieved by only 10% of participants in the PItch-Only and Combined conditions, and only 20% in the Time-Only condition.

**Table 4.** Performance across deciles for each test

|  | 1 | 2 | 3 | 4 | 5 | 6 | 7 | 8 | 9 | 10 |
| --- | --- | --- | --- | --- | --- | --- | --- | --- | --- | --- |
| Focus, pitch | 0.48 | 0.57 | 0.67 | 0.79 | 0.88 | 0.93 | 0.98 | 0.98 | 1.00 | 1.00 |
| Focus, time | 0.43 | 0.53 | 0.60 | 0.66 | 0.71 | 0.76 | 0.79 | 0.82 | 0.85 | 0.92 |
| Focus, both | 0.45 | 0.56 | 0.67 | 0.86 | 0.94 | 0.96 | 0.98 | 0.98 | 1.00 | 1.00 |
| Phrase, pitch | 0.45 | 0.53 | 0.60 | 0.65 | 0.69 | 0.73 | 0.77 | 0.81 | 0.85 | 0.89 |
| Phrase, time | 0.52 | 0.58 | 0.63 | 0.67 | 0.74 | 0.79 | 0.84 | 0.86 | 0.88 | 0.93 |
| Phrase, both | 0.47 | 0.56 | 0.68 | 0.75 | 0.81 | 0.86 | 0.88 | 0.91 | 0.96 | 0.98 |

**Figure 2.**
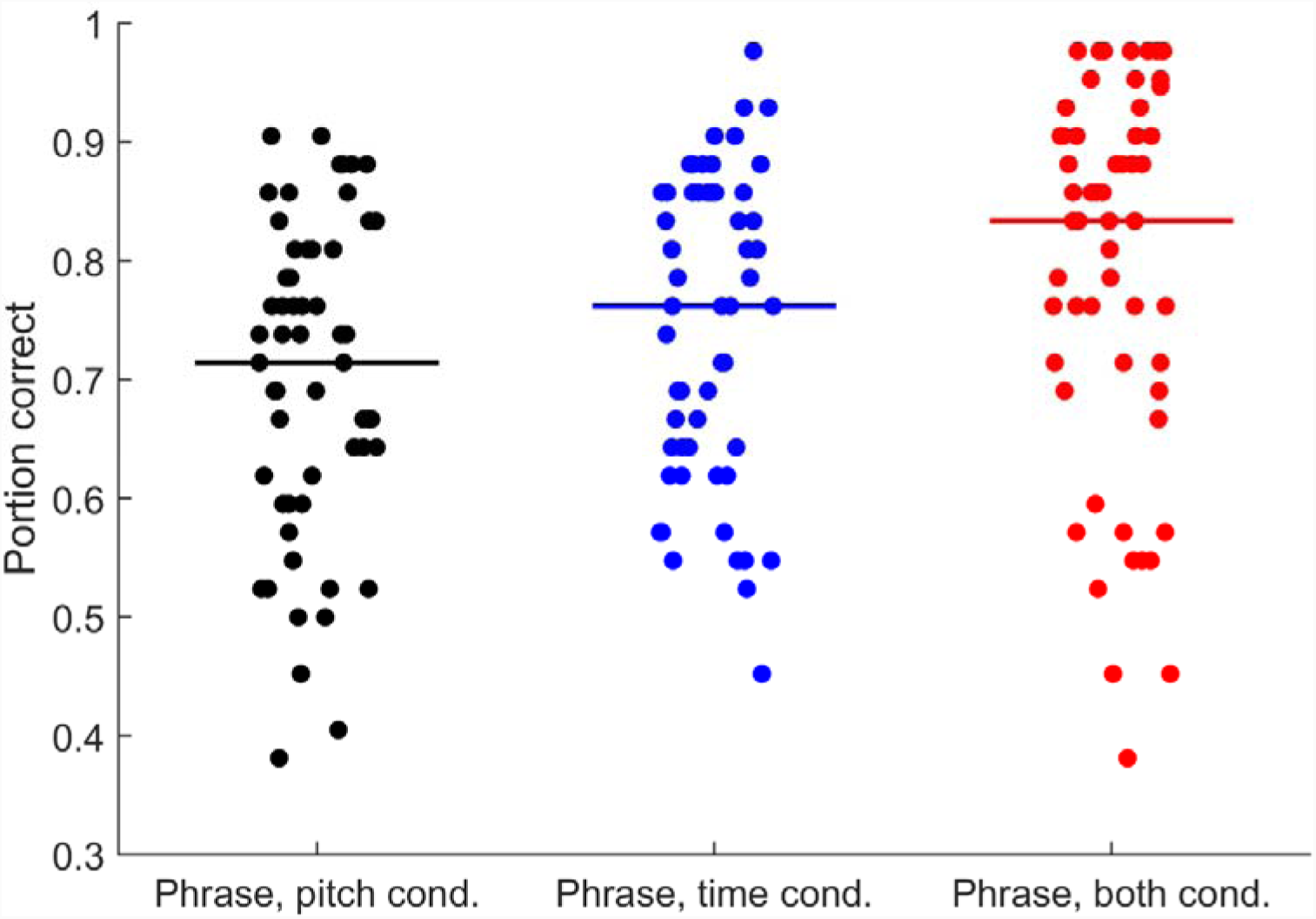
Performance across all 57 participants in each condition of the phrase perception test. Horizontal lines indicate median performance.

**Figure 3.**
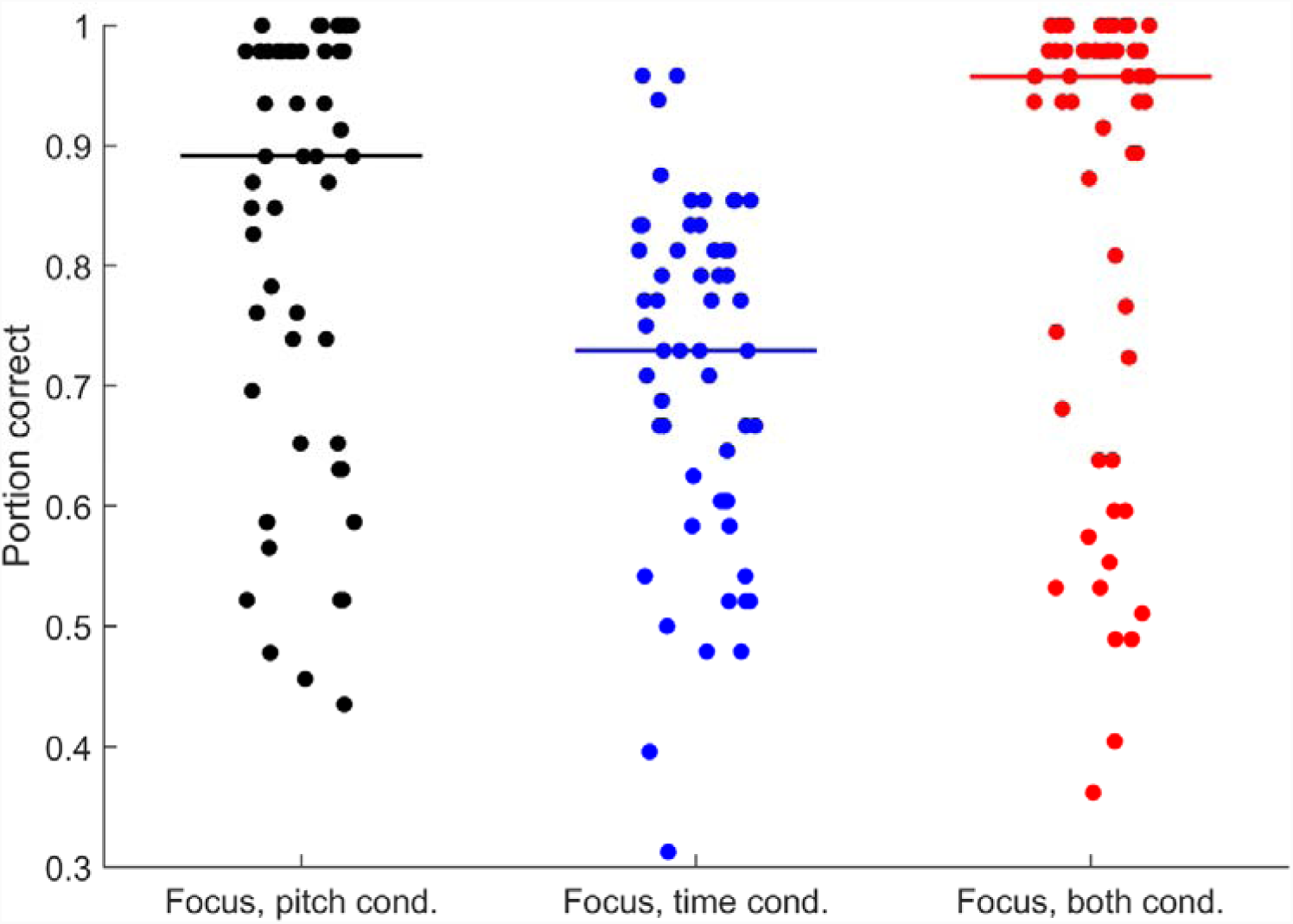
Performance across all 57 participants in each condition of the focus perception test. Horizontal lines indicate median performance.

### 3.2 Subtest reliability

Cronbach’s alpha was used to calculate reliability for each of the six subtests. For the focus tests, reliability was 0.90 for the pitch condition, 0.80 for the time condition, and 0.92 for the both condition. For the phrase tests, reliability was 0.77 for the pitch condition, 0.75 for the time condition, and 0.87 for the both condition. To summarize, reliability tended to be highest for the both condition, and reliability was somewhat higher for the focus tests than for the phrase tests. Overall, however, these reliability scores compare favourably with those of other batteries of prosody perception (Kalathottukaren, Purdy, & Ballard, 2015).

### 3.3 Comparison between conditions

To examine the relative usefulness of pitch and time cues in the perception of phrase boundaries and linguistic focus we conducted a 2 × 3 repeated measures ANOVA with test (phrase versus focus) and condition (both, pitch, and time) as factors. There was a main effect of test (F(1,56) = 22.45, p < 0.001), indicating that participants performed better on the focus test than the phrase test. There was also a main effect of condition (F(2,112) = 47.12, p < 0.001) and an interaction between test and condition (F(2,112) = 58.83, p < 0.001). Bonferroni-corrected post-hoc paired t-tests revealed that for focus perception, participants performed better on the Combined condition compared to the Time-Only condition (t(56) = 9.93, p < 0.001) but not compared to the Pitch-Only condition (t(56) = 1.62, p > 0.1). Moreover, participants performed better on the Pitch-Only condition compared to the Time-Only condition (t(56) = 8.11, p < 0.001). For phrase perception, there was a main effect of condition (F(2, 112) = 26.7, p < 0.001). Bonferroni-corrected post-hoc paired t-tests revealed that participants performed better on the both condition compared to both the Pitch-Only (t(56) = 7.52, p < 0.001) and Time-Only (t(56) = 4.09, p < 0.001) conditions. Moreover, participants performed better on the Time-Only condition relative to the Pitch-Only condition (t(56) = 3.14, p < 0.01). These results suggest that for focus perception, pitch was a more useful cue than time, while for phrase perception, time was a more useful cue than pitch. Moreover, across both focus and phrase perception, the presence of an additional cue was generally useful to listeners.

### 3.4 Relationships between conditions

Pearson’s correlations were used to examine the relationship between performance across all six subtests. Correlations are listed in Table 5, and relationships between all six variables are displayed in scatterplots in Figure 4. False Discovery Rate (Benjamini & Hochberg (1995) procedure) was used to correct for multiple comparisons. Correlations between all conditions were significant, but varied in strength. Generally correlations between subtests within each prosody test were stronger than correlations between prosody tests. For example, the correlation between performance in the pitch condition and time condition of the focus perception test was r = 0.70, while the correlation between performance in the pitch condition of the phrase test and the time condition of the focus perception test was r = 0.46.

**Table 5.**
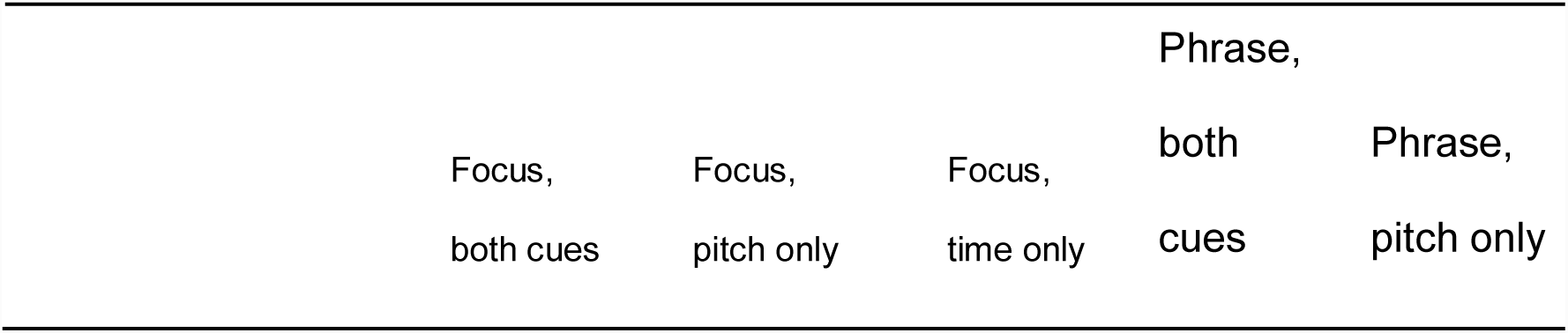
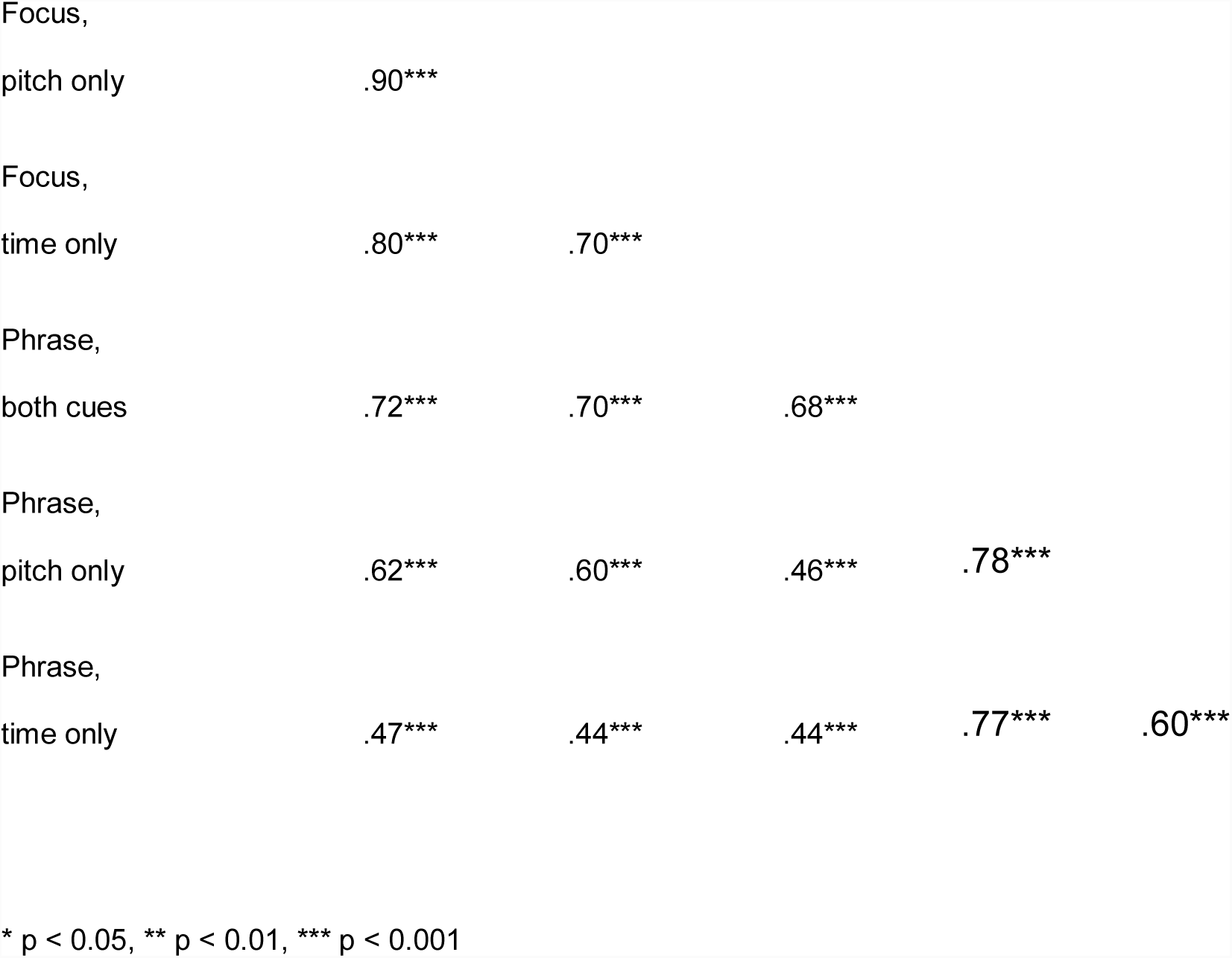
Pearson’s correlations between performance on all six prosody perception sub-tests.

**Figure 4.**
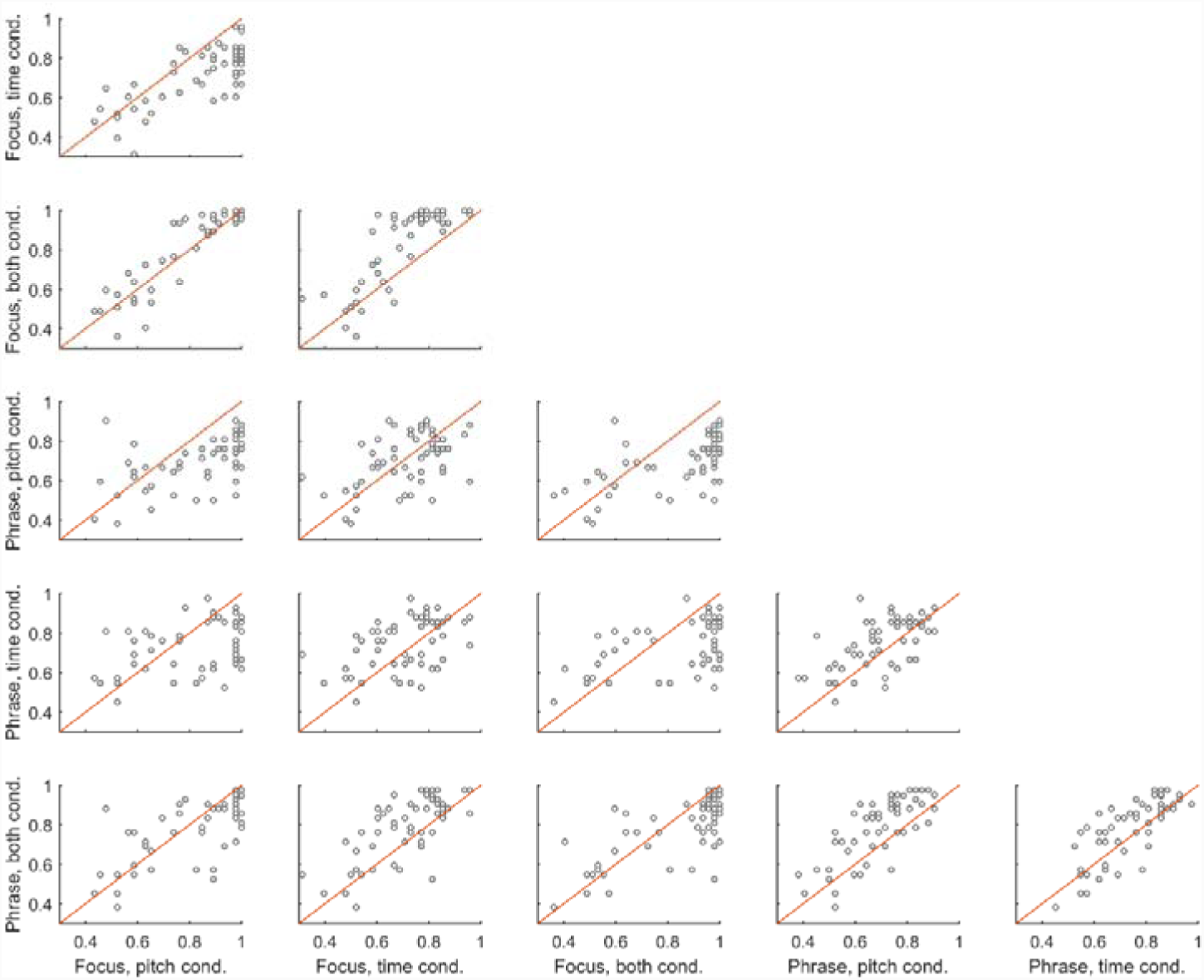
Scatterplots displaying the relationship between performance across each possible pair of all six conditions. The diagonal red line indicates the identity line.

The correlation data does not indicate that subtests requiring analysis of similar perceptual cues correlate more strongly. For example, the correlation between the two time conditions is no stronger than the correlation between the time condition of the focus test and the pitch condition of the phrase test. This result raises the question of whether the pitch and time conditions are, indeed, indexing different aspects of prosody perception. We investigated this question by conducting two multiple linear regressions, one for focus perception and one for phrase perception, with performance on the pitch and time conditions as independent variables and performance on the both condition as the dependent variable. For focus perception, we found that pitch performance (standardized β = 0.67, p < 0.001) and time performance (standardized β = 0.33, p < 0.001) explained independent variance in performance in the both cues condition. This suggests that perception of focus draws on both pitch and duration perception, but that pitch is relatively more important. For phrase perception, we also found that pitch performance (standardized β = 0.50, p < 0.001) and time performance (standardized β = 0.48, p < 0.001) explained independent variance in performance in the both cues condition. This suggests that perception of phrase boundaries draws on both pitch and duration perception, and that both cues are relatively equally important.

## 4. Discussion

Here we have presented a new battery of prosody perception which is suitable for examining prosody perception in adults. This instrument could facilitate investigation of a number of research questions, such as whether difficulties with prosody perception in individuals with dyslexia or ASD extend into adulthood. This battery could also be used to test the hypothesis that musical training can enhance focus and phrase boundary perception. This possibility is supported by findings that musical training is linked to enhanced encoding of the pitch of speech (Bidelman, Gandour, & Krishnan, 2009; Marques, Moreno, & Luís Castro, 2007; Moreno & Besson, 2005; Musacchia, Sams, Skoe, & Kraus, 2007; Wong, Skoe, Russo, Dees, & Kraus, 2007) and syllable durations (Chober, Marie, Francois, Schon, & Besson, 2011) and that musicians are better than nonmusicians at detecting stress contrasts (Kolinsky et al., 2009) and discriminating statements from questions based on intonational contours (Zioga, Luft, & Bhattacharya, 2016).

### 4.1 Adaptive difficulty

The test stimuli for the Multidimensional Battery of Prosody Perception (MBOPP) were created using speech morphing software. As a result, the test difficulty is fully customizable (because researchers can select the stimuli with desired cue magnitude) without compromising ecological validity and naturalisticalness of the stimuli. The data reported here were collected by setting prosodic cue size to medium levels. This resulted in data that largely avoided both floor and ceiling effects in typically developing adults, although there was some evidence of ceiling performance in the Pitch-Only and both cues conditions of the focus perception test. This suggests that to equate difficulty across the focus and phrase perception tests the cue size for the focus perception test should be slightly lower than that for the phrase perception test.

Given that cue size was set here at 50% of maximum, there remains quite a bit of scope for lowering the difficulty of the test in order to make it appropriate for other populations who may have lower prosody perception skills such as children, or adults with perceptual difficulties. The ability to modify cue size on a fine-grained level also enables researchers to modify test difficulty on an item-by-item basis. This could have two important uses. First, adaptive prosody perception tests could allow researchers to rapidly find participants’ thresholds for accurate prosody perception by modifying test difficulty in response to participants’ performance, enabling the use of shorter test protocols. And second, adaptive prosody perception training paradigms could be created by ensuring that participants are presented with stimuli at a difficulty level that is neither so easy as to be trivial nor so difficult as to be frustrating.

### 4.2 Independent modification of individual cues

Another novel feature of the MBOPP is the ability to modify the size of pitch and duration cues independently. This makes possible investigations into whether prosody perception deficits are cue-specific in certain populations. For example, we have demonstrated using the MBOPP that adults with amusia demonstrate impaired focus perception in the Pitch-Only condition, but perform similarly to typically developing adults on the Time-Only condition (Jasmin et al., 2018). Investigating the cue specificity of prosody perception deficits is one way to test the hypothesis that difficulties with prosody perception in a given population stem from auditory deficits. For example, some individuals with ASD have difficulty perceiving prosodic cues to phrase boundaries (Diehl et al., 2008) and linguistic focus (Peppé et al. 2011). ASD has also been linked to impaired duration discrimination (Brenner et al., 2015; Karaminis et al., 2016; Martin, Poirier, & Bowler, 2010) but preserved pitch discrimination and memory for pitch sequences (Heaton, Hudry, Ludlow, & Hill, 2008; Jarvinen-Pasley, Wallace, Ramus, Happé, & Heaton, 2008; Stanutz, Wapnick, & Burack, 2014). If prosodic deficits in ASD stem from abnormalities in auditory processing, then they should reflect the unique auditory processing profile of individuals with ASD, and prosodic impairments should be greater for perception and production of duration-based prosodic cues compared to pitch-based prosodic cues.

On the other hand, if impairments are present across all conditions, regardless of the acoustic cue presented, this would suggest that prosodic difficulties in ASD stem primarily from modality-general deficits in the understanding of emotional and pragmatic aspects of language.

### 4.3 The role of pitch and durational cues in focus and phrase perception

Our results suggest that pitch and duration cues play somewhat different roles in focus perception versus phrase perception. Specifically, performance on the Pitch-Only condition surpassed performance on the Time-Only condition for focus perception, while performance on the Time-Only condition surpassed performance on the Pitch-Only condition for phrase perception. This finding is consistent with the literature on the acoustic correlates of prosody, as pitch changes have been shown to be a more reliable cue to linguistic focus than syllabic lengthening (Breen et al., 2010). On the other hand, durational cues such as preboundary lengthening and increased pauses have been shown to be more reliable cues to phrase structure than pitch changes (Choi et al., 2005; but see Streeter, 1978, who showed that pitch and durational cues are used to a roughly equal extent by listeners). This suggests that impairments in pitch versus duration perception, which have been shown to be dissociable (Hyde & Peretz, 2004; Kidd et al., 2007), may not have equal effects on different aspects of prosody perception. For example, individuals with impaired pitch perception may have greater difficulties with the perception of linguistic focus than with phrase boundary perception, as we have demonstrated in amusics (Jasmin et al., 2018).

Speech tends to be structurally redundant, i.e. a given speech category is often conveyed by multiple acoustic cues simultaneously. This property may make speech robust to both external background noise (Winter, 2014) and internal “noise” related to imprecise representation of auditory information (Patel, 2014). In support of this idea, we found that performance on the both cues condition surpassed that of either single-cue condition for phase perception, in alignment with previous findings that rising pitch and increased duration are more effective cues to phrase boundaries when presented simultaneously (Cumming, 2010). On the other hand, performance on the both cues condition of the focus perception test did not exceed that of the Pitch-Only condition. This suggests that different prosodic features may vary in the extent to which they are conveyed by redundant cues and, therefore, the extent to which they are vulnerable to the degradation of a particular cue, either due to external or internal noise.

### 4.4 Limitations

The MBOPP currently has a number of limitations which should be kept in mind by users but could be addressed in future versions of the battery. First, all test items were spoken by a single talker. As a result, the relative usefulness of pitch versus duration cues for a given prosodic feature may reflect that talker’s idiosyncratic patterns of cue use rather than, more generally, the usefulness of those cues across talkers. Second, only English test items are included, limiting the extent to which the battery can be generalized to other populations. And third, currently only two aspects of prosody perception are included, focus perception and phrase boundary detection. Stress perception and emotion perception are two particularly important aspects of prosody perception which will be included in future versions.

## Acknowledgments

The work was funded by a Wellcome Trust Seed Award #109719/Z/15/Z to A.T.T., a Reg and Molly Buck Award from SEMPRE to K.J., and a Leverhulme Trust Early Career Fellowship to K.J.

## Open practices statement

The MBOPP auditory materials used in these experiments are available at http://researchdata.bbk.ac.uk/id/eprint/37

